# Polyphosphate uses mTOR, pyrophosphate, and Rho GTPase components to potentiate bacterial survival in *Dictyostelium*

**DOI:** 10.1101/2023.05.03.539237

**Authors:** Ryan J. Rahman, Ramesh Rijal, Shiyu Jing, Te-An Chen, Issam Ismail, Richard H. Gomer

## Abstract

Human macrophages and the eukaryotic microbe *Dictyostelium discoideum* ingest bacteria by phagocytosis, and then kill the ingested bacteria. Some pathogenic bacteria secrete linear chains of phosphate residues (polyphosphate; polyP), and the polyP causes the phagocytes to not kill the ingested bacteria. In *D. discoideum*, the effect of polyP requires the G protein-coupled receptor GrlD, suggesting that polyP uses a signal transduction pathway to inhibit killing of ingested bacteria. Here we show that in addition to GrlD, the *D. discoideum* polyP signaling pathway requires the GPCR interacting arrestin-like protein AdcB, inositol hexakisphosphate kinase A (I6kA), the Rho GTPase RacE, and the TOR component Lst8. *D. discoideum* also secretes polyP, and at high concentrations polyP inhibits *D. discoideum* cytokinesis. The polyP inhibition of bacterial killing pathway does not appear to involve many of the polyP inhibition of cytokinesis pathway components. These data suggest the intriguing possibility that if there is a similar polyP inhibition of bacterial killing pathway in macrophages, pharmacologically blocking this pathway could potentiate macrophage killing of pathogenic bacteria.

**Importance:** Although most bacteria are quickly killed after phagocytosis by a eukaryotic cell, some pathogenic bacteria prevent their killing after phagocytosis. Pathogenic *Mycobacterium* species secrete polyP, and the polyP is necessary for the bacteria to prevent their killing after phagocytosis. Conversely, exogenous polyP prevents the killing of ingested bacteria that are normally killed after phagocytosis by human macrophages and the eukaryotic microbe *Dictyostelium discoideum*. This suggests the possibility that in these cells, a signal transduction pathway is used to sense polyP and prevent killing of ingested bacteria. In this report, we identify key components of the polyP signal transduction pathway in *D. discoideum*. In cells lacking these components, polyP is unable to inhibit killing of ingested bacteria. The pathway components have orthologues in human cells, and an exciting possibility is that pharmacologically blocking this pathway in human macrophages would cause them to kill ingested pathogens such as *M. tuberculosis*.

## Introduction

Polyphosphate (polyP) is a linear polymer of three to hundreds of orthophosphate residues, and is found in all kingdoms of life (1). PolyP is predominantly synthesized by polyphosphate kinase 1 (Ppk1), which is highly conserved in more than 100 prokaryotes, including more than 20 major bacterial pathogens and a few eukaryotes such as *Dictyostelium discoideum* (2). Bacteria lacking Ppk1 lose essential functions for survival, such as cell motility, biofilm formation, and pathogenicity (3–5). The pivotal role polyP plays in pathogenesis has marked Ppk1, which is absent in humans, as a potential target to block pathogenicity (6).

In mammalian cells, polyP (n∼50-150 phosphates) is in platelet and mast cell granules and is released during injury or cell activation to potentiate blood clotting cascades (7). Platelet-released polyP also triggers release of neutrophil extracellular traps (8), induces macrophage differentiation and neutrophil chemoattraction during wound healing (9), but inhibits peritoneal macrophage chemotaxis to the sites of infection and tissue damage (10). PolyP can also signal through receptors such as P2Y1 and RAGE to potentiate pro-inflammatory responses of endothelial cells as well as mediate communication among astrocytes (11, 12).

Macrophages fight bacterial infections by phagocytosis (13). During phagocytosis, a bacterium is engulfed into a phagosome, which then acidifies and fuses with a lysosome to form a phagolysosome to kill and digest the ingested bacterium (14). Pathogens such as *Mycobacterium tuberculosis* (*Mtb*) prevent their killing in human macrophages by inhibiting phagosome acidification and fusion of the phagosome with the lysosome (15). We previously observed that *M. smegmatis* and *Mtb* secrete extracellular polyP (16), that exogenous polyP inhibits phagosome acidification and lysosome activity, and that polyP potentiates the survival of non-pathogenic *Escherichia coli* in human macrophages (16). Other workers found that exogenous polyP potentiates pathogenic *E. coli* survival in a murine model of sepsis (17). Conversely, treatment of human macrophage and *Mtb* co-cultures with the polyP degrading enzyme exopolyphosphatase (PPX), or reduced expression of Ppk1 in *M. smegmatis*, reduced the survival of these bacteria in human macrophages (16, 18). Together, these results suggest that extracellular polyP produced by some pathogenic bacteria contributes to their survival in macrophages (16).

*D. discoideum* is a eukaryotic microbe that primarily feeds on bacteria by phagocytosis (19–21). Many *D. discoideum* proteins involved in phagocytosis are conserved in human neutrophils and macrophages (22). Proliferating *D. discoideum* cells accumulate extracellular polyP, and as the cell density increases to near the point where the cells are about to overgrow their food supply and starve, the concomitant high levels of extracellular polyP inhibit cytokinesis, but not the accumulation of cell mass so that the cells will be large and have high nutrient reserves when they starve (23). The extracellular polyP is sensed by the putative G protein-coupled polyP receptor GrlD (24). PolyP inhibits proliferation through distinct mechanisms based on nutrient availability,as GrlD partially mediates this effect in high-nutrient conditions while GrlD and a small GTPase RasC are necessary for the effect in low-nutrient conditions (24). In addition to GrlD and RasC in low-nutrient conditions, polyP requires the G protein component Gβ, the Ras guanine nucleotide exchange factor GefA, phosphatase and tensin homolog (PTEN), phospholipase C (PLC), inositol 1,4,5-trisphosphate (IP3) receptor-like protein A (IplA), polyphosphate kinase 1 (Ppk1), and the TOR complex 2 component PiaA to inhibit proliferation (25). With the exception of *grlD^−^*, *rasC^−^*, and *piaĀ*, the strains had reduced but non-zero responses to polyP, suggesting the existence of parallel pathways mediating polyP effects (25).

In *D. discoideum*, polyP acts via the polyP receptor GrlD to potentiate *E. coli* survival (16). We previously observed that *E. coli*, which do not accumulate detectable extracellular polyP get killed after phagocytosis by *D. discoideum*, while *M. smegmatis*that accumulate detectable extracellular polyP survive better after phagocytosis than *E. coli* (16). As in macrophages, reduced expression of *ppk1* in *M. smegmatis* bacteria reduces their survival in *D. discoideum*, and the addition of exogenous polyP potentiates the survival (16). Together, this suggests the intriguing possibility that there is a signal transduction pathway whereby either extracellular polyP or polyP secreted by a bacterium in a phagosome induces cells to not fuse the phagosome with a lysosome. In this report, we screened *D. discoideum* mutants to elucidate polyP signal transduction pathways that are needed for polyP to potentiate the survival of ingested bacteria in *D. discoideum*. We find that extracellular polyP requires the GPCR interacting arrestin-like protein AdcB, inositol hexakisphosphate kinase A (I6kA), the Rho GTPase RacE, and the TOR component Lst8 to potentiate *E. coli* survival in *D. discoideum*.

## Results

### PolyP requires a G protein-coupled receptor, but does not require G-protein subunits, to potentiate the survival of *Escherichia coli* in *D. discoideum*

Wild-type (WT) *D. discoideum* cells accumulate extracellular polyP, and the polyP concentrations (≥ 470 μg/mL) corresponding to high cell densities (≥ 1 x 10^7^ cells/ ml) inhibit proliferation, macropinocytosis, exocytosis, and killing of ingested bacteria (23, 26). PolyP concentrations between 5 and 47 μg/mL, which do not affect proliferation, macropinocytosis, or exocytosis, inhibit the killing of ingested bacteria in *D. discoideum* (16). To identify components of the polyP signal transduction pathway that potentiate the survival of bacteria in *D. discoideum*, 37 available mutants derived from 7 parental strains were tested for sensitivity to polyP-mediated *E. coli* survival as previously described (16) (Table 1). In these assays, *Dictyostelium* cells are allowed to ingest *E. coli* bacteria, the uningested bacteria are washed off, any remaining uningested bacteria are killed with the antibiotic gentamicin, which does not kill ingested bacteria (27), and at 4 and 48 hours, aliquots of the *Dictyostelium* cells are lysed with a detergent that does not kill *E. coli*, and the ingested bacteria are plated to obtain a count of live bacteria. Adding extracellular polyP has little effect on the bacterial survival at 4 hours, and strongly potentiates survival at 48 hours (16). To determine if polyP-mediated changes in intracellular *E. coli* numbers correspond to altered ingestion or digestion, the efficiency of phagocytic engulfment of Zymosan A bioparticles was measured for all strains. The data are graphed in nine groups: parental/wild-type cells (Figures 1A, **2A, S1A, and S2A**), polyP receptor, G-proteins, and arrestin-like proteins (Figures 1B, **2B, S1B, and S2B**), proteins involved in polyP production (Figures 1C, **2C, S1C, and S2C**), GTPases (Figures 1D, **2D, S1D, and S2D**), phospholipase C (PLC)/IP3 pathway components (Figures 1E, **2E, S1E, and S2E**), PI3 kinase signal transduction pathway components (Figures 1F, **2F, S1F, and S2F**), TOR complex components/protein kinases (Figures 1G, **2G, S1G, and S2G**), autophagy pathway components (Figures 1H, **2H, S1H, and S2H**), and cytoskeleton regulating proteins (Figures 1I, **2I, S1I, and S2I**). At 4 hours, polyP did not affect the number of viable ingested *E. coli* in the parental/WT strains or any of the mutants, with the exception of cells lacking the arrestin-like protein AdcC, where polyP caused a slight drop in the number of viable internalized E. coli (**Figures S1A-I** and Table 1). At 48 hours, polyP potentiated the viability of ingested *E. coli* in all of the parental strains (Figure 1A and Table 1). In addition to there being differences (in the absence of exogenous polyP) between these parental strains in the number of viable bacteria at 4 and at 48 hours (Figures 1A and **S1A** and Table 1), there were also differences in phagocytosis (Figures 2A and **S2A** and Table 1), indicating some strain-dependent differences in phagocytosis and phagosome/lysosome physiology.

**Figure 1:**
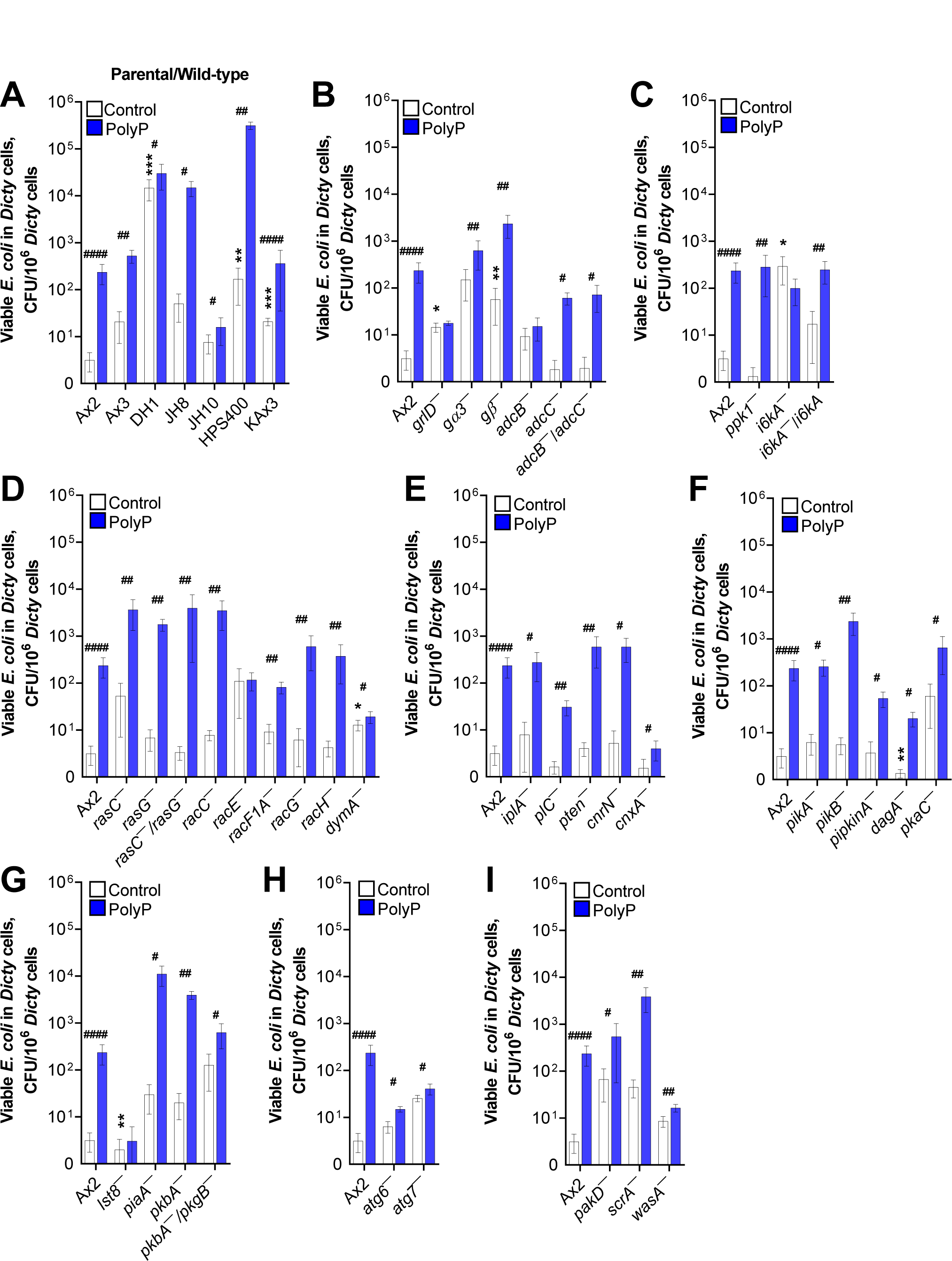
PolyP potentiates the long-term survival of ingested *E. coli* in *D. discoideum*. (A-I) *D. discoideum* (Dicty) cells were incubated with *E. coli*, uningested *E. coli* were removed, and the number of viable ingested *E. coli* per 10^6^ *D. discoideum* cells in the absence (Control) or the presence of added polyphosphate (PolyP) was determined at 48 hours. Values are mean ± SEM from 5 independent experiments for each mutant/parental strain and 16 independent experiments for Ax2 wild type. * p < 0.05, ** p < 0.01, and *** p < 0.001 by Mann-Whitney test comparing the indicated mutant to its parental strain (Table 1), or the indicated parental strain to Ax2, in the absence of added polyP. # p < 0.05, ## p < 0.01, and #### p < 0.0001 by Mann-Whitney test comparing control to polyP for the indicated strain.

**Figure 2:**
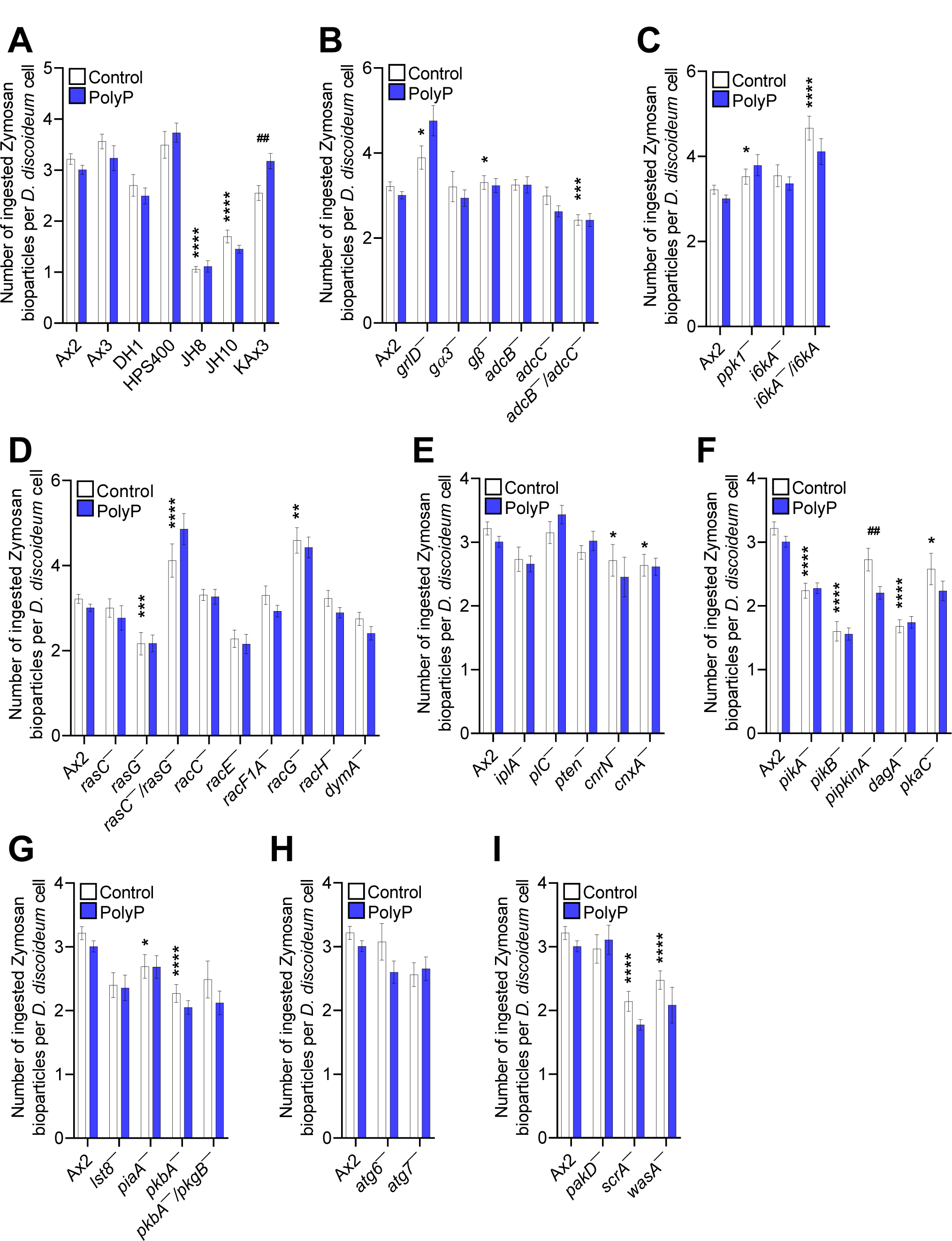
PolyP has negligible effect on the uptake of Zymosan A bioparticles in most strains. (A-I) The number of Zymosan A bioparticles ingested per *D. discoideum* cell in 60 minutes in the presence or absence of 15 µg/ml polyP was determined. Values are mean ± SEM from 3 independent experiments. * p < 0.05, ** p < 0.01, *** p < 0.001, and **** p < 0.0001 comparing the indicated mutant to its parental strain, or the indicated parental strain to Ax2, in the absence of added polyP by Mann-Whitney test. ## p <0.01 comparing control and polyP for the indicated strain by Mann-Whitney test.

**Table 1:**
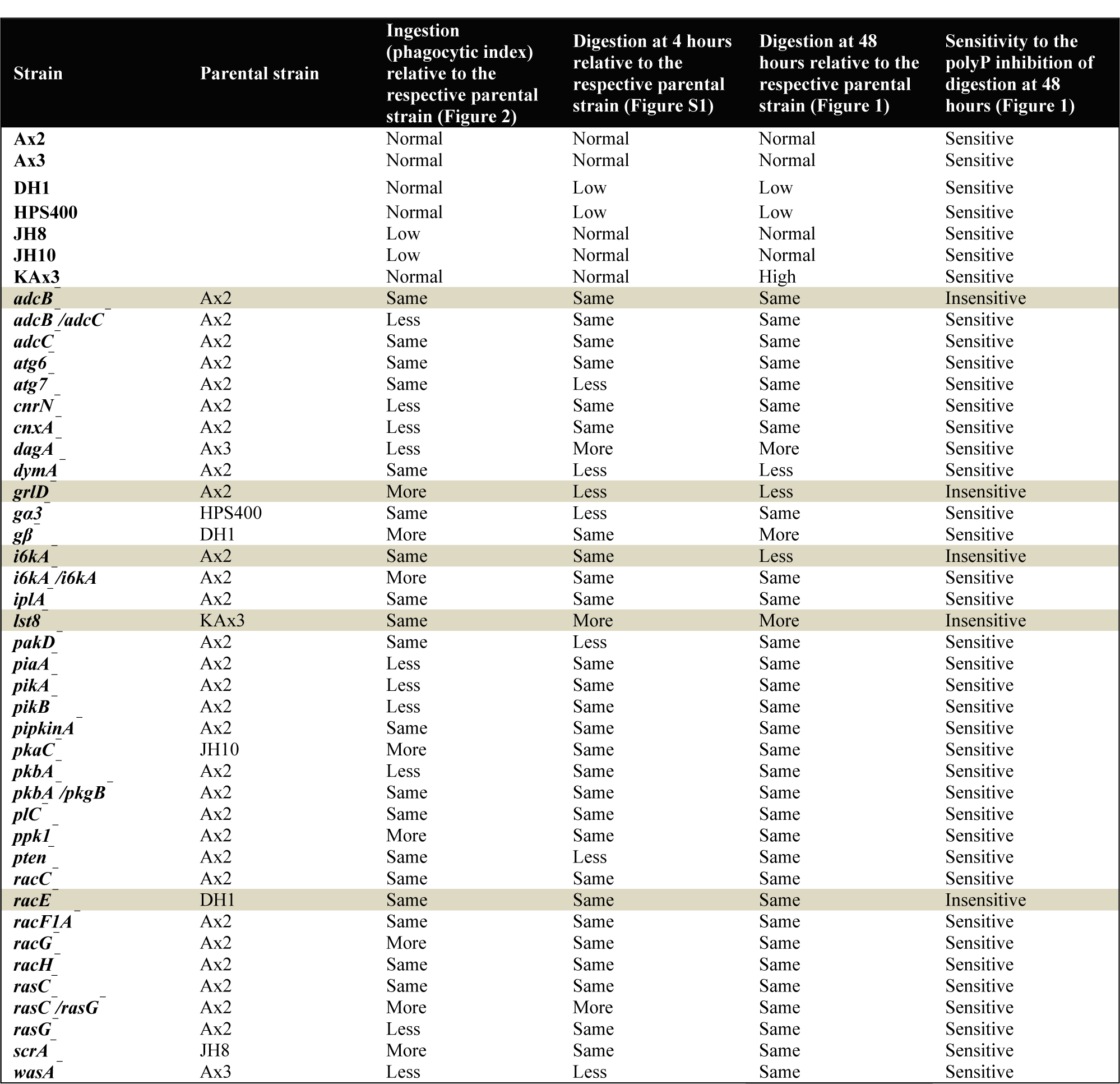
*D. discoideum* sensitivity to polyP. Forty-four *D. discoideum* strains including seven parental wild-type strains were tested for sensitivity to polyP for ingestion of zymosan bioparticles and digestion of ingested *E. coli* at 4 and 48 hours. Parental strains were compared against Ax2, which was considered normal, and mutants were compared against their parental strains and considered less, same, or more for the indicated parameters tested. Inherent decrease or increase in digestion of ingested bacteria in mutants were indicated as more or less than the respective parental strains. Mutants that responded or did not respond to polyP-mediated inhibition of the digestion of ingested bacteria at 48 hours were considered sensitive or insensitive, respectively.

Cells lacking the putative G protein-coupled polyP receptor GrlD are insensitive to polyP-induced proliferation inhibition (24, 25). Cells lacking the heterotrimeric G protein subunits Gβ and Gα3 have a reduced but non-zero sensitivity to polyP (25), suggesting the existence of an additional pathway downstream of GrlD. GrlD, but not Gβ and Gα3, were needed for polyP to potentiate *E. coli* viability at 48 hours (Figure 1B and Table 1), suggesting that polyP uses GrlD to activate a downstream pathway that bypasses, or partially bypasses G proteins.

### PolyP requires the arrestin domain containing protein AdcB to potentiate *E. coli* survival

Persistent activation of GPCRs is dampened by phosphorylation of the cytoplasmic region of the receptors (28). Arrestins are scaffolding proteins that bind to the carboxyl terminus of phosphorylated GPCRs, uncouple GPCRs from their cognate G proteins, and turn off G protein mediated signaling (29, 30). In addition to desensitizing GPCRs, arrestins can also function as adaptor proteins for GPCR trafficking and G protein-independent signaling (31, 32). *D. discoideum* does not possess true arrestins, but has genes encoding six arrestin domain containing proteins (AdcA-F) (33). AdcB but not AdcC was needed for polyP potentiation of bacterial survival at 48 hours, although polyP potentiated bacterial survival in cells lacking both arrestin-like proteins (Figure 1B and Table 1), suggesting the possibility that AdcB mediates polyP signaling downstream of the GrlD receptor, and that lack of both AdcB and AdcC might cause upregulation of a yet unknown component, possibly one of the four other arrestin-like proteins, that can compensate for the lack of AdcB.

### PolyP requires an inositol hexakisphosphate kinase to potentiate *E. coli* survival

Polyphosphate kinase 1 (Ppk1) synthesizes polyP from ATP (2, 34). *D. discoideum* cells lacking Ppk1 (*ppk1^−^*) possess undetectable levels of intracellular polyP (35), and reduced but detectable levels of extracellular polyP (23). Inositol hexakisphosphate kinase (IP6K) synthesizes inositol pyrophosphates IP7 and IP8 from IP6 (36). *D. discoideum* cells lacking the IP6K homolog I6kA accumulate reduced levels of extracellular polyP (23). At 48 hours, polyP increased the number of viable ingested *E. coli* in *ppk1^−^* cells but not *i6kĀ* cells, and the latter defect was rescued in *i6kĀ/i6kA* cells (Figure 1C and Table 1), indicating that *D. discoideum* requires I6kA to mediate polyP induced survival of ingested *E. coli*.

### PolyP requires the RhoGTPase RacE to potentiate *E. coli* survival

Ras and Rho GTPases are involved in variety of cellular processes including proliferation, differentiation, cell motility, cell polarity and trafficking of vesicles and macromolecules (37, 38). PolyP requires RasC to inhibit proliferation and induce development of *D. discoideum* cells (25, 39), whereas the Rho GTPase RacC was not necessary for polyP mediated proliferation inhibition (25). The ability of polyP to potentiate the survival of ingested bacteria at 48 hours did not require RasC, RasG, RacC, RacF1A, RacG, RacH, or the large GTPase dynamin, which is involved in membrane remodeling during endocytosis and phagocytosis (40, 41) (Figure 1D and Table 1). However, the polyP effect did require the Rho GTPase RacE (42) (Figure 1D and Table 1).

### PolyP does not require several PLC/IP3 pathway components to potentiate *E. coli* survival

Phospholipase C, which converts PIP2 to diacylglycerol and inositol 1,4,5-trisphosphate (IP3), and the IP3 receptor-like protein IplA, are required for polyP mediated *D. discoideum* proliferation inhibition (25, 43). Phosphatase and TENsin homolog (PTEN) and the PTEN-like phosphatase CnrN catalyze the conversion of PIP3 to PIP2 (44, 45), and are involved in many cellular processes including proliferation and cell migration (25, 44, 45). PTEN but not CnrN is involved in polyP mediated *D. discoideum* proliferation inhibition (25). Calnexin (Cnx) is a calcium binding protein and interacts with IP3 receptors (46). Other potential components of PIP3-associated pathways include phosphatidylinositol kinases PikA and PikB, phosphatidylinositol phosphate kinase A (*pipkinĀ*), the pleckstrin homology (PH) domain containing and PIP3 binding cytosolic regulator of adenylyl cyclase protein CRAC (DagA), and the cAMP-dependent protein kinase A catalytic subunit PkaC. Poly potentiated bacterial survival at 48 hours in *iplĀ*, *plC^−^*, *pten^−^*, *cnrN^−^*, *cnxĀ*, *pikĀ, pikB^−^, pipkinĀ dagĀ*, and *pkaC^−^* cells (Figures 1E and **F** and Table 1), suggesting that polyP does not use these components of the PLC/IP3 pathway to potentiate the survival of ingested bacteria.

### PolyP requires the TOR complex protein Lst8 to potentiate *E. coli* survival

The Target of Rapamycin (TOR) forms two distinct signaling complexes, TOR complex 1 (TORC1) and TORC2 (47). In mammals, TORC1 activation promotes anabolic metabolism and blocks catabolic processes such as autophagy and lysosome biogenesis (48, 49). TORC2 is involved in cytoskeletal reorganization during chemotactic cell movement (50–52). TORC1 and TORC2 complexes have shared and unique components. TOR and Lst8 are present in both TORC1 and TORC2 complexes, whereas PiaA (mammalian Rictor) is unique to TORC2 (47). PolyP requires PiaA to inhibit proliferation of *D. discoideum* cells, whereas cells lacking Lst8 (*lst8^−^*) are sensitive to polyP mediated proliferation inhibition (25). TORC2 regulates the activity of protein kinase B (PKB) (53). PolyP required Lst8 but not PiaA, the Akt/PKB protein kinase PkbA, or PkbA and the SGK family protein kinase PkgB to potentiate bacterial survival (Figure 1G and Table 1). Together, these data suggest that polyP might regulate TOR signaling via Lst8 to potentiate bacterial survival.

### PolyP does not require the autophagy proteins Atg6 and Atg7 to potentiate *E. coli* survival

Autophagy is used to degrade and recycle cytoplasmic materials in eukaryotic cells (54), and TOR signaling regulates autophagy (55). *D. discoideum* cells feed on bacteria to acquire nutrients in the natural environment. The cells use autophagy machinery to kill bacteria only when the bacteria escape the phagosome in a process called xenophagy (56). *D. discoideum* uses the autophagy proteins Atg6 and Atg7 for autophagosome formation (57). PolyP potentiated the survival of ingested bacteria in *atg6^−^* and *atg7^−^* cells, however, the effect of polyP on *atg6^−^* and *atg7^−^* was mild compared to WT cells. (Figure 1H and Table 1). Together, these data suggest that polyP may not require components of the autophagy pathway to potentiate bacterial survival.

### PolyP does not require selected cytoskeletal proteins to potentiate *E. coli* survival

Actin and actin associated proteins play critical roles during phagocytic uptake and early phagosome formation processes (58–60). Although polyP concentrations (705 µg/ml) corresponding to very high cell densities reduced levels of actin cytoskeleton proteins (39), the lower polyP concentration that potentiates bacteria survival in WT cells did not require the cytoskeleton-associated proteins p21-activated kinase D (PakD), or Wiskott Aldrich Syndrome protein family proteins SCAR or WasA to potentiate bacterial survival (Figure 1I) suggesting that polyP prevents the killing of ingested bacteria without requiring these proteins.

### Defective polyP sensitivity is not due to a defect in the parental strain or defective phagocytosis

Cells lacking GrlD, AdcB, I6kA, RacE, and Lst8 (with genotypes verified by PCR, **Figure S3**) do not potentiate bacterial survival at 48 hours in response to polyP. Ax2 is the parental strain of *grlD^−^*, *adcB^−^*, and *i6kĀ*, DH1 is the parental strain of *racĒ*, and KAx3 is the parental strain of *lst8^−^* (Table 1). All of these parental strains increased bacterial survival in response to polyP (Figure 1A and Table 1), indicating that the defects in the above mutants are not due to a defect in the parental strain. Phagocytosis, as measured by the number of ingested zymosan particles, was normal in most of the above strains with defective polyP responses, with the exception of *grlD^−^*, which had somewhat higher phagocytosis (Figure 2 and Table 1). Compared to the respective parental strains, there was no consistent effect of these mutations on the percent of cells ingesting beads. Cells lacking GrlD were normal, a slightly higher percentage of cells lacking AdcB and I6kA ingested beads, and a somewhat lower percentage of cells lacking RacE and Lst8 ingested beads (**Figure S2**). PolyP had no significant effect on these percentages (**Figure S2**). Together, these results indicate that the above proteins are part of a mechanism where extracellular polyP potentiates the survival of ingested bacteria after phagocytosis of the bacteria.

### PolyP inhibits proteasome activity

Cells label proteins with ubiquitin to induce their degradation (61). To determine if polyP potentiation of bacterial survival grossly affects ubiquitinated protein levels, Western blots of *D. discoideum* lysates from the 4 and 48 hour bacterial survival assays were stained with anti-ubiquitin antibodies. PolyP slightly increased the level of ubiquitinated proteins at 4 hours compared to control, but ubiquitinated protein levels were similar in control and polyP treated cells at 48 hours (Figure 3A and **B**). Together, these data suggest that polyP may transiently increase ubiquitinated protein levels, that this effect on ubiquitination may be compensated after 4 hours by some unknown mechanism, and that this short-term effect of polyP may be independent of the effect of polyP on facilitating the long term survival of bacteria.

**Figure 3:**
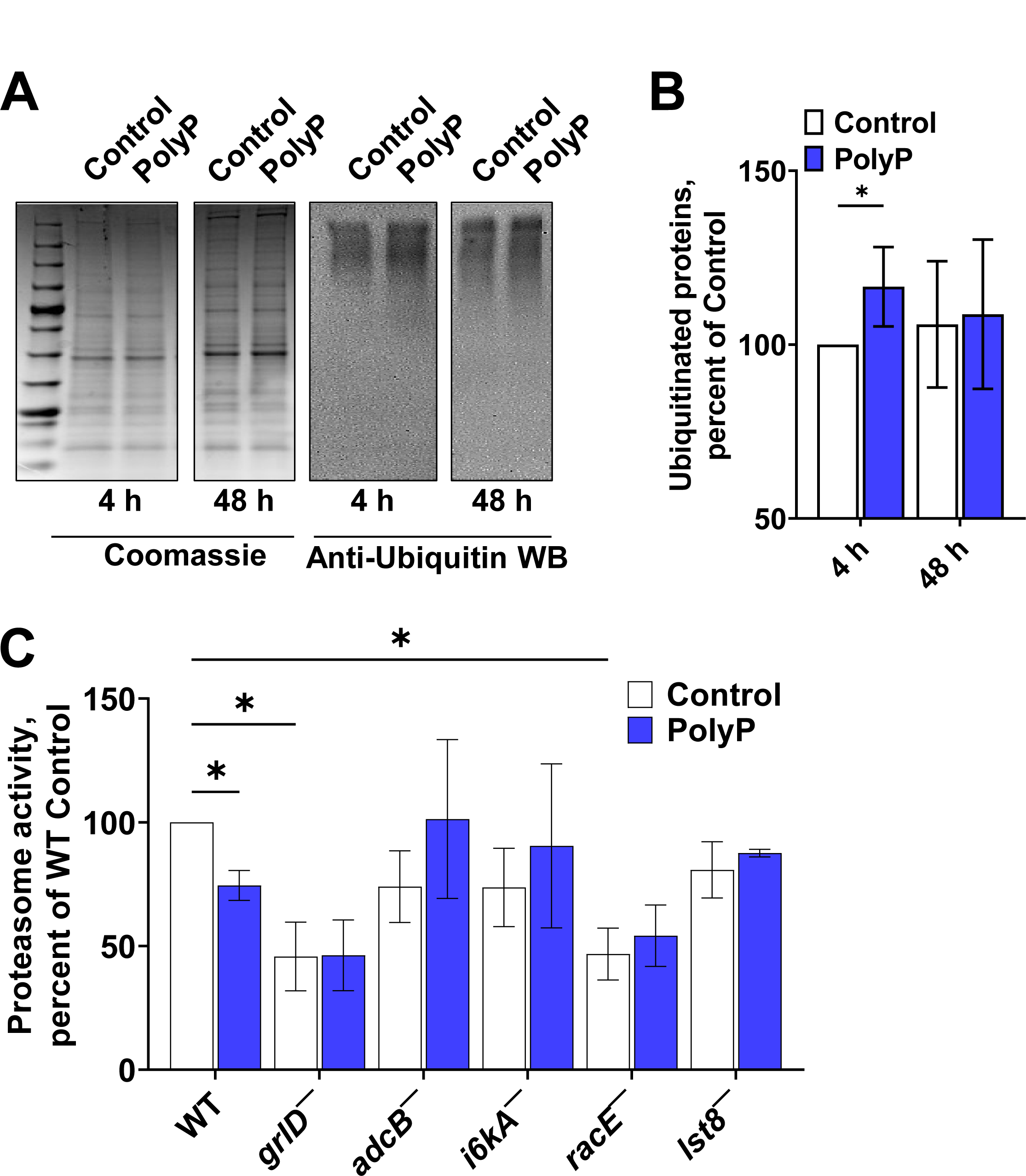
PolyP affects protein ubiquitination and proteasome activity. (A) Coomassie stains and anti-ubiquitin western blot images for 4 and 48 hour Ax2 lysates in the presence and absence of polyP. Images are representative of 3 independent experiments. (B) The integrated staining intensities in the anti-ubiquitin western blots were quantified as a percent of the 4-hour control. (C) Proteasome activity of Ax2 (WT) and the indicates strains in the presence or absence of 15 µg/ml polyP were measured and normalized to the Ax2 controls. Values are mean ± SEM from 3 independent experiments. * p < 0.05 by Mann-Whitney test.

We previously found that 705 μg/ml polyP (the extracellular polyP concentration in stationary phase cultures) inhibits proteasome activity in *D. discoideum* (39). PolyP requires GrlD and RasC to inhibit proteasome activity in all nutrient conditions (39), but to inhibit proliferation, polyP inhibits proteasome activation only in low nutrient conditions, indicating that polyP inhibits proliferation of *D. discoideum* using different pathways depending on the nutritional conditions (39). To determine if the relatively low concentrations of polyP that potentiate the survival of *E. coli* also inhibit proteasome activity, we tested the effect of 15 μg/ml polyP on proteasome activity in Ax2 and the mutants that are insensitive to polyP-induced bacterial survival. PolyP reduced proteasome activity in Ax2 cells, and although *grlD^−^* and *racĒ* cells had reduced basal proteasomal activity, polyP did not significantly affect proteasome activity in *grlD^−^, adcB^−^, i6kĀ, racĒ*, or *lst8^−^* cells (Figure 3C). This indicates that GrlD, AdcB, I6kA, RacE, and Lst8 may be parts of a polyP signaling pathway that reduces proteasome activity, although whether this is associated with, or independent of, the effects of this same pathway on survival of ingested bacteria remains unknown.

### PolyP requires AdcB and Lst8 to inhibit cell proliferation

We previously observed that 705 μg/ml polyP inhibits the proliferation of WT *D. discoideum* cells (23), and that the loss of GrlD, I6kA or TORC2 complex protein PiaA reduces the ability of polyP to inhibit proliferation in a low nutrient (25% HL5) medium (25). To determine if the proteins that mediate polyP potentiation of bacterial survival also affect polyP inhibition of proliferation in normal nutrient conditions, we examined the effect of 705 μg/ml polyP on proliferation of mutants in SIH and HL5 media. In SIH, compared to WT cells, polyP had a reduced ability to inhibit the proliferation of *grlD*^−^ and *lst8*^−^ cells, and in HL5, polyP had a reduced ability to inhibit the proliferation of *grlD*^−^, *adcB*^−^, and *lst8*^−^ cells (Figure 4). Together, this indicates that some but not all polyP potentiation of bacterial survival signal transduction pathway components also mediate polyP proliferation inhibition, suggesting a partially shared pathway.

**Figure 4:**
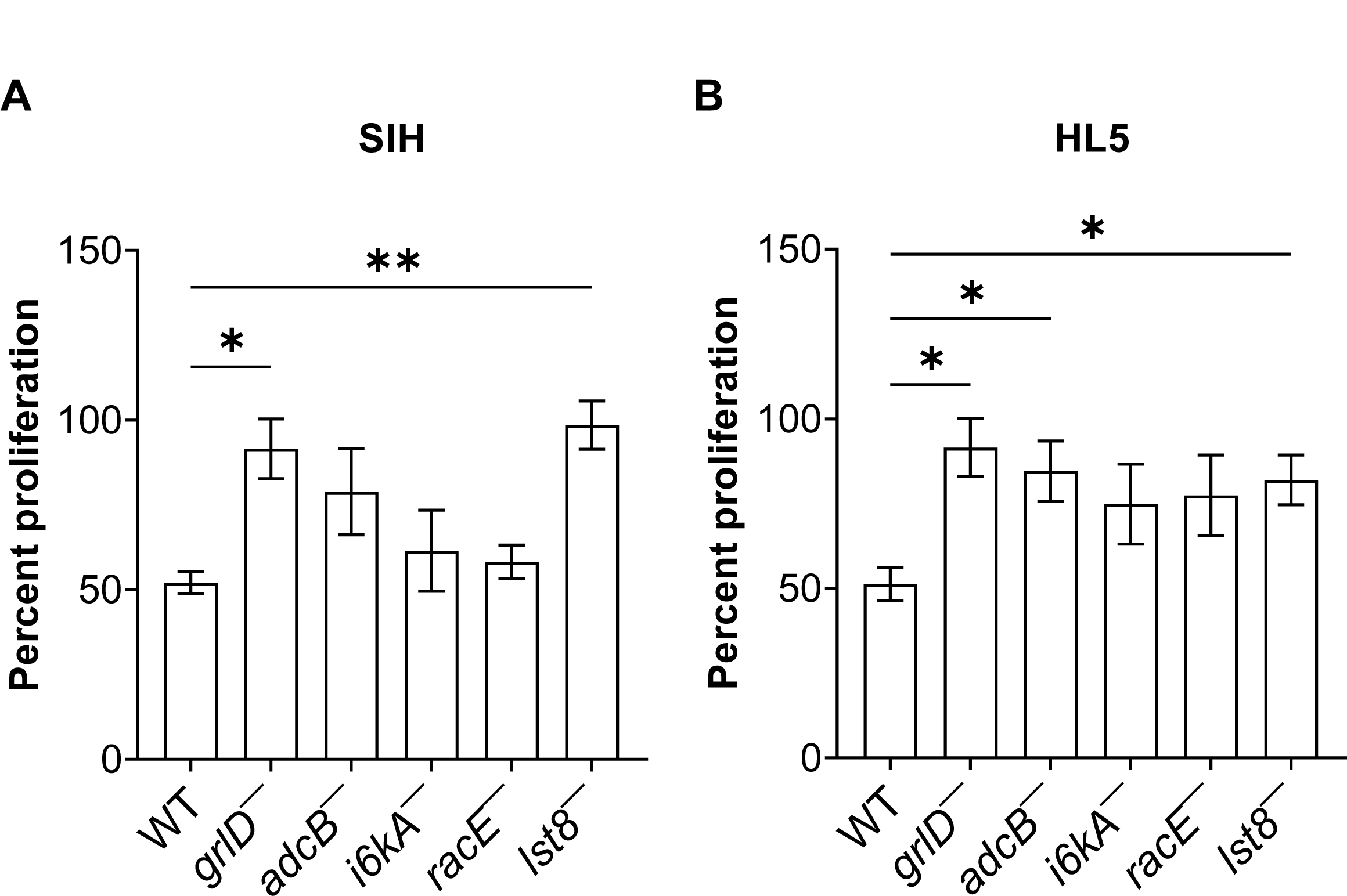
PolyP requires GrlD, AdcB, and Lst8 to inhibit the proliferation of *D. discoideum* cells. Cells were cultured in either SIH (A) or HL5 (B) with or without 705 µg/ml polyP for 24 hours. The increase in cell density in the presence of polyP was calculated as a percentage of the increase in cell density in the absence of polyP for each strain. Values are mean ± SEM from 3 independent experiments. * p < 0.05 and ** p < 0.01 by Mann-Whitney test.

## Discussion

In this report, we found that to inhibit the killing of ingested bacteria in *D. discoideum*, polyP requires, in addition to the G protein-coupled receptor GrlD (16, 39), AdcB, I6kA, RacE, and Lst8. PolyP also appears to use AdcB and Lst8, but not I6kA and RacE, to inhibit proliferation. However, to inhibit killing of ingested bacteria, polyP does not require other components of the signal transduction pathway that it uses to inhibit cell proliferation such as Gβ, RasC, PakD, PiaA, PTEN, PLC, IplA, Ppk1, PiaA, and PkaC (23, 25), suggesting that polyP uses partially overlapping signal transduction pathways to inhibit proliferation and the killing of ingested bacteria (Figure 5).

**Figure 5:**
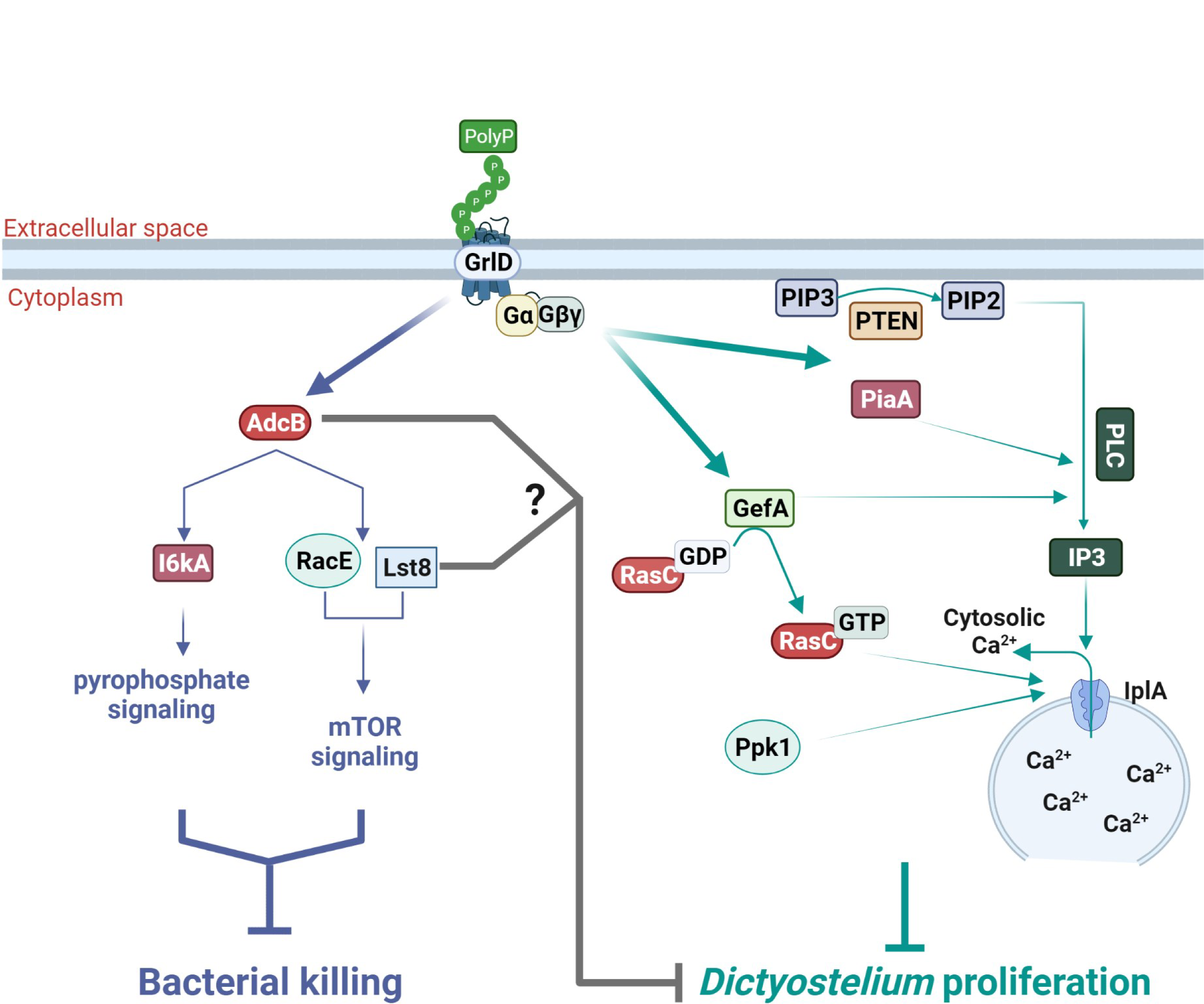
Hypothesized polyP signal transduction pathways that inhibit the killing of bacteria in *D. discoideum* and pathways that inhibit cell proliferation. PolyP binds to the GrlD receptor, and the polyP signal is relayed downstream of GrlD by the arrestin-like protein AdcB, inositol hexakisphosphate kinase A (I6kA), the Rho GTPase RacE, and the TOR component Lst8 to inhibit the killing of ingested bacteria. AdcB and Lst8 also help polyP to inhibit *D. discoideum* proliferation. To inhibit proliferation, polyP signaling downstream of GrlD receptor is also mediated by Gαs and Gβγ, PTEN, and PLC, which cleaves PIP2 to IP3 and DAG. When IP3 binds to the IP3 receptor IplA, Ca^2+^ is released into the cytosol. The increase of IP3 and cytosolic Ca^2+^ levels by polyP depends on PTEN, PLC, and IplA. GefA converts GDP-bound RasC to GTP-bound RasC. Both GefA and PiaA are essential for the upregulation of IP3 levels by polyP, while RasC and Ppk1 are necessary for the increase of cytosolic Ca^2+^ levels. The proposed pathway diagram was created using BioRender.com.

Although polyP requires the G protein subunit Gβ to inhibit proliferation in 25% HL5 (25), polyP does not require Gα3 and Gβ to potentiate the survival of ingested bacteria in *D. discoideum*, but rather uses the arrestin-like protein AdcB downstream of GrlD. Arrestins are scaffolding proteins that deactivate a G protein-coupled receptor by binding to the cytoplasmic domain of the receptor, which induces a conformational change that allows the release of the G proteins subunits bound to the receptor (29, 30). Therefore, polyP appears to activate different pathways immediately downstream of GrlD to inhibit proliferation and the killing of ingested bacteria.

Eukaryotic cells including *D. discoideum* utilize autophagy for intracellular degradation of cytoplasm and organelles (57, 62-66). In addition to preventing phagosome-lysosome fusion, pathogens such as *M. tuberculosis* and *Salmonella typhimurium* break out of phagosomes and enter the cytosol (67). The eukaryotic cell then degrades the disrupted phagosome and attempts to kill the pathogen in the cytosol by autophagy (67–69). *S. typhimurium*, which requires polyphosphate kinase to prevent their killing in human macrophages (70), show increased survival in *D. discoideum* cells lacking the autophagy proteins Atg6 and Atg7 (64). However, polyP does not need Atg6 and Atg7 to increase the survival of *E. coli* in *D. discoideum* cells. Assuming that *E. coli* are unable to break out of the phagosome, this indicates that polyP may not require autophagy machinery to potentiate the survival of ingested bacteria in phagosomes, and that autophagy is a second line of defense against pathogens that do escape the phagosome.

The high polyP concentrations that inhibit proliferation of *D. discoideum* cells also inhibit proteasome activity in *D. discoideum* and human leukemia cell lines (39), and inhibition of proteasome activity has been suggested as a potential therapeutic for cancer (71). The low polyP concentration (15 µg/ml) that does not inhibit proliferation but potentiates survival of bacteria inhibits proteasome activity in WT cells but not in *grlD^−^*, *adcB^−^ i6kĀ, lst8^−^, and racĒ* cells. This indicates that components of the polyP pathway which inhibits killing of ingested bacteria also inhibit proteasome activity. One possibility is that this pathway allows some *D. discoideum* cells to sense the relatively low concentrations of extracellular polyP, and from this sense that they are near each other and will eventually overgrow their food supply and begin to conserve energy by storing nutrients (not killing some of ingested bacteria) and decreasing protein degradation.

In this report, we found that polyP inhibits killing of ingested bacteria in *D. discoideum* cells using signal transduction pathway components and mechanisms that have orthologues in human cells. PolyP from ingested pathogenic bacteria inhibits their killing in human macrophages (16). An intriguing possibility is that macrophages have a pathway similar to the *Dictyostelium* pathway to sense polyP, and that blocking this pathway could induce macrophages to kill internalized pathogens such as *M. tuberculosis*.

### Contact for Reagent and Resource Sharing

Further information and requests for reagents may be directed to, and will be fulfilled by, the authors Ramesh Rijal and Richard Gomer.

## Materials and Methods

### D. discoideum cell culture

*D. discoideum* strains were obtained from the *Dictyostelium* Stock Center (72) and were Ax2 (DBS0237699), Ax3 (DBS0235542), KAx3 (DBS0266758), DH1 (DBS0235700), JH8 (DBS0236454), JH10 (DBS0236449), and HPS400 (DBS0236312), *adcB*^−^ (DBS0350443)*, adcC*^−^ (DBS0350646), *adcB*^−^*/adcC*^−^ (DBS0350445), *atg6*^−^ (DBS0236344), *atg7*^−^ (DBS0236372), *cnrN*^−^ (DBS0302655), *cnxA*^−^ (DBS0236189), *dagA*^−^ (DBS0235559), *dymA*^−^ (DBS0347874), *grlD*^−^ (DBS0350227)*, gβ*^−^ (DBS0236531), *gα3*^−^ (DBS0235986)*, iplA*^−^ (DBS0236260), *i6kA*^−^ (DBS0236426), *i6kA*^−^*/i6kA* (23), *lst8*^−^ (DBS0236517)*, pakD*^−^ (DBS0350281), *piaA*^−^ (DBS0349879) *, pikA*^−^ (DBS0350197), *pikB*^−^ (DBS0350198), *pipkinA*^−^ (DBS0236779), *pkaC*^−^ (DBS0236783)*, pkbA*^−^ (DBS0349876), *pkbA*^−^/*pkgB*^−^ (DBS0236785), *plC*^−^ (DBS0267124), *ppk1*^−^ (DBS0350686), *pten*^−^ (DBS0236830), *racC*^−^ (DBS0350272), *racE*^−^ (DBS0235413), *racF1*^−^ (DBS0351505), *racG*^−^ (DBS0236849), *racH*^−^ (DBS0236850), *rasC*^−^ (DBS0236853), *rasG*^−^ (DBS0236862), *rasC*^−^*/rasG*^−^ (DBS0236858), *scrA*^−^ (DBS0236926) and *wasA*^−^ (*wasA*^−^ strain was a kind gift from Robert Insall, Beatson Institute for Cancer Research, Glasgow, UK) (2, 42, 45, 51, 52, 72-103). *D. discoideum* cell cultures were maintained at 21 °C in type 353003 100 mm tissue culture dishes (Corning, Durham, NC) in 10 ml of SIH defined minimal medium (Formedium, Norfolk, England) and were selected with appropriate antibiotics and supplements as previously described (16, 52).

### Polyphosphate preparation

Sodium polyphosphate (polyP) of average chain length of 45 monomers (16) (Cat#S0169, Spectrum, New Brunswick, NJ) was used for all assays. PolyP stock solutions were prepared as previously described (16).

### Bacterial survival assay and phagocytosis

*E. coli* K12 survival in *D. discoideum* was performed as previously described (16). Phagocytic index is a measurement of the uptake of particles by phagocytes (104). Fluorescence microscopy was used to visualize ingested Alexa 594-labeled Zymosan-A yeast BioParticles (Cat#Z23374, Thermo Fisher Scientific) in *D. discoideum* as described in (16).

### Genotype verification

The genotype of *D. discoideum* strains that were insensitive to polyP (*grlD*^−^, *adcB*^−^, *i6kA*^−^, *lst8*^−^, and *racE*^−^) were verified by PCR as previously described (52) using the specific primer pairs listed in Table 2.

**Table 2:**
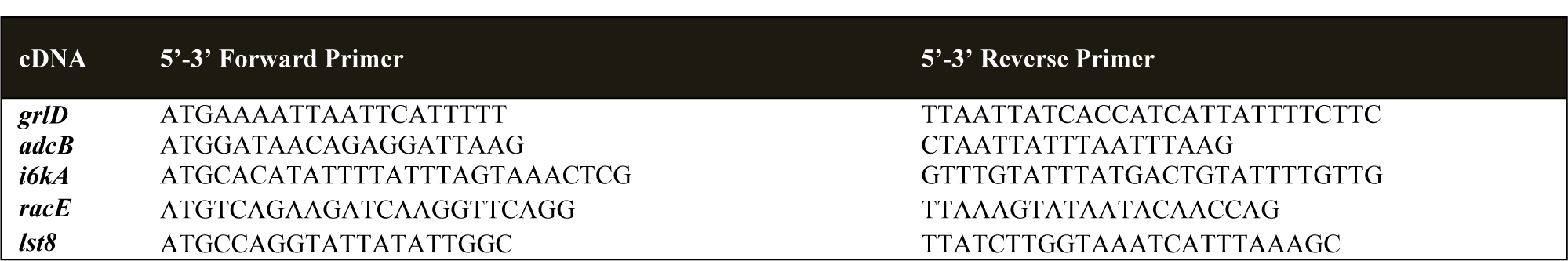
Oligonucleotides for genotyping of polyP insensitive mutants by PCR.

### Immunoblotting

1 ml of 10^6^ *D. discoideum* cells in SIH were seeded in a type 353047 24-well tissue culture plate (Corning), and either 15 µg/ml polyP from a 15 mg/ml stock in PBM (20 mM KH_2_PO_4_, 1 mM MgCl_2_, 0.01 mM CaCl_2_, pH adjusted to 6.1 with KOH) or an equivalent volume of PBM was added. At 4 or 48 hours, the 24-well plate was centrifuged at 500 x g for 3 minutes, and the supernatant was replaced by 1 ml of PBM. This step was repeated once. The 24-well plate was centrifuged at 500 x g for 3 minutes, the supernatant was discarded, and cells at the bottom of each well were lysed with 100 µl of 1X SDS sample buffer. The lysates were collected, and western blots and Coomassie staining of gels was performed as previously described (52). Western blots were stained with 1:1000 diluted mouse monoclonal anti-ubiquitin antibodies (Cat# 3936T; Cell Signaling Technology) to detect ubiquitinated proteins.

### Proteasome activity assay

100 µl of *D. discoideum* cells in SIH at 10^6^ cells/ ml were seeded in type 353219, 96-well, black/clear, tissue-culture-treated, glass-bottom plates (Corning), spun down at 500 x g for 3 minutes, the medium was changed to SIH or SIH containing 15 µg/ml polyP, and the 96 well plate was incubated in a Tupperware container with wet paper towels (for humidity) for 48 hours. Proteasome activity was measured using a proteasome activity kit (#MAK172, Sigma, St Louis, MO) following the manufacturer’s instructions.

### Proliferation inhibition

Proliferation of *D. discoideum* strains in the presence or absence of 705 µg/ml polyP was measured as previously described (23).

### Statistical analysis

Statistical analyses were performed using Prism 9 (GraphPad, San Diego, CA) and the tests indicated in the figure legends. A p < 0.05 was considered to be significant.

## Acknowledgements

We thank Dr. Robert Insall, Beatson Institute for Cancer Research, Glasgow, UK for the gift of *wasA*^−^ cells, the *Dictyostelium* stock center for other cells.

## Funding

Ryan Rahman was supported by awards from the Arnold and Mabel Beckman Foundation, the Goldwater Scholar Foundation, and the Astronaut Scholar Foundation. This work was supported by National Institutes of Health grants GM118355 and GM139486.

## Author Contributions

R. R. designed and performed experiments, analyzed data, and wrote the paper. R. J. R. performed experiments, analyzed data, and wrote the paper, S. J. performed experiments, T. C. performed experiments, and R. H. G. coordinated the study, revised the paper, and acquired funding.

## Declaration of Interests

No competing interests declared.

